# Macrophage Function is Regulated by NPM1-mediated 2’-O-methylation

**DOI:** 10.1101/2020.04.18.048223

**Authors:** Paolo Sportoletti, Daphna Nachmani, Luisa Riccardi, Arati Khanna-Gupta, Jian Chen, Andrea Marra, Nancy Berliner, John G. Clohessy, Pier Paolo Pandolfi

**Affiliations:** Cancer Research Institute, Beth Israel Deaconess Medical Center, Harvard Medical School, Boston, MA, 02115, USA; Institute of Hematology-Centro di Ricerca Emato-Oncologica (CREO), University of Perugia, Italy; Division of Hematology, Brigham and Women’s Hospital, Harvard Medical School, Boston, MA, 02115, USA; MBC, Department of Molecular Biotechnology and Health Sciences, University of Torino, Torino, TO, 10126, Italy.

## Abstract

The *NPM1* gene is frequently a target of genetic alteration in hematological tumors, particularly of the myeloid lineage. Complete inactivation of *Npm1* in the mouse disrupts primitive hematopoiesis and results in embryonic lethality. *Npm1* heterozygosity produces features similar to those of MDS that progress to overt leukemia, and specific point mutations of *Npm1* lead to bone marrow failure due to loss of hematopoietic stem cells. However, little is known about NPM1’s role in mature, differentiated cells. Here we generated a conditional mouse mutant to inactivate *Npm1* across the myelomonocytic lineage, and investigated its ability to influence macrophage maturation and function. We found that *Npm1* is not required to maintain macrophage viability, while its loss in mature macrophages reduces production of reactive oxygen species, chemotactic properties and phagocytic capacity. Taking advantage of our recently established *Npm1*^*D180del*^ mouse model of ribosome dysfunction and hematological disease, we identify cellular translation and rRNA 2’-O-methlyation as a crucial element in controlling macrophage function. These analyses demonstrate a role for *Npm1* in adult immune cells, and reveal the importance of translation regulation in macrophage function.

**Statement of significance:** Macrophages are a major component of the immune response to various insults including to cancer. Here we show that NPM1, the most frequently mutated gene in acute myeloid leukemia, displays a critical role in macrophage function, and we identify ribosome deregulation as one of the underlying mechanisms.

## Introduction

NPM1 is an abundant phosphoprotein with multiple cellular functions (*1*), and which is essential for hematopoietic development. Complete loss of *Npm1* results in early embryonic lethality characterized by defective primitive hematopoiesis (*2*). Conditional complete inactivation in the adult mouse hematopoietic system results in bone marrow failure characterized by dysmegakaryopoiesis, defective erythroid maturation, dysplastic and low platelet counts, and dysplastic neutrophils; all features of ribosome dysfunction syndromes (*3*). The human *NPM1* gene is frequently found to be altered in different myeloid tumors including *NPM1-mutated* acute myeloid leukemia (AML) (*4, 5*), acute promyelocytic leukemia (APL) and MDS (*6*). Indeed, *Npm1*^+*/-*^ mice display features resembling some of those identified in human myelodysplastic syndrome (MDS) and an increased susceptibility to develop myeloid malignancies (*2, 5*). Recently, we have demonstrated that NPM1 fine-tunes cellular translation, by regulating rRNA modifications involved in maintaining ribosomal integrity. We show that specific mutations in *NPM1* lead to altered rRNA modifications and hence ribosome dysfunction, resulting in bone marrow failure and expansion of the myeloid compartment(*3*). This myeloid-bias implies that NPM1 contributes myeloid development at multiple stages and not only at the stem cell level. However, compelling dissection of NPM1’s functional role across the myeloid lineage is still fragmentary.

Our analysis demonstrates that *Npm1* loss affects macrophage effector functions and suggests impairment in the myelomonocytic transcriptional hub in mice. Mechanistically, we show these affects are in part mediated through ribosome regulation by NPM1. These findings reveal NPM1 and the ribosome as key determinants of immune function.

## Results

### MRP8-Cre mice induce NPM1 loss in macrophages

In order to establish the role of *Npm1* across the myelomonocytic lineage beyond the hematopoietic stem cells we initiated our study by crossing our mice carrying a floxed conditional allele of *Npm1 (Npm1*^*fl/fl*^) (*3*) with the *MRP8-Cre* transgenic mouse model. In these mice the Cre recombinase protein is expressed under the control of the myeloid specific *MRP8* promoter (*7*) enabling recombination at the common myeloid progenitor (CPM) compartment. The *Npm1*^*fl/fl*^;*MRP8-Cre*^+^ mice generated were found to be viable and were born in expected Mendelian ratios (data not shown).

In order to assess the efficiency of *Npm1* deletion by Cre recombinase we first analyzed mature myeloid cells within the bone marrow by sorting for Gr1^+^;Mac1^+^ cells from both *Npm1*^*fl/fl*^;*MRP8-Cre*^+^ and *Npm1*^*fl/fl*^;*MRP8-Cre*^*-*^ mice. Consistent with the MRP8 expression pattern, recombination of the floxed *Npm1* allele was observed in Gr1^+^;Mac^+^ positive myeloid cells in the bone marrow of *Npm1*^*fl/fl*^;*MRP8-Cre*^+^ mice, while the *Npm1* locus was unaffected in *Npm1*^*fl/fl*^;*MRP8-Cre*^*-*^ (Figure 1A). Importantly, no recombination of the floxed *Npm1* allele was observed in B220^+^ sorted bone marrow cells from *Npm1*^*fl/fl*^;*MRP8-Cre*^+^ mice. However, immunofluorescence analysis of bone marrow cells clearly revealed the presence of the NPM1 protein present in doughnut shaped granulocytes in *Npm1*^*fl/fl*^;*MRP8-Cre*^+^ (Figure 1B). The presence of the NPM1 protein in granulocytes was also confirmed by western blot analysis of Gr1^+^;Mac1^+^ sorted cells where the expression was comparable in *Npm1*^*fl/fl*^;*MRP8-Cre*^+^ and *Npm1*^*fl/fl*^;*MRP8-Cre*^*-*^ controls (Figure 1C).

**Figure 1.**
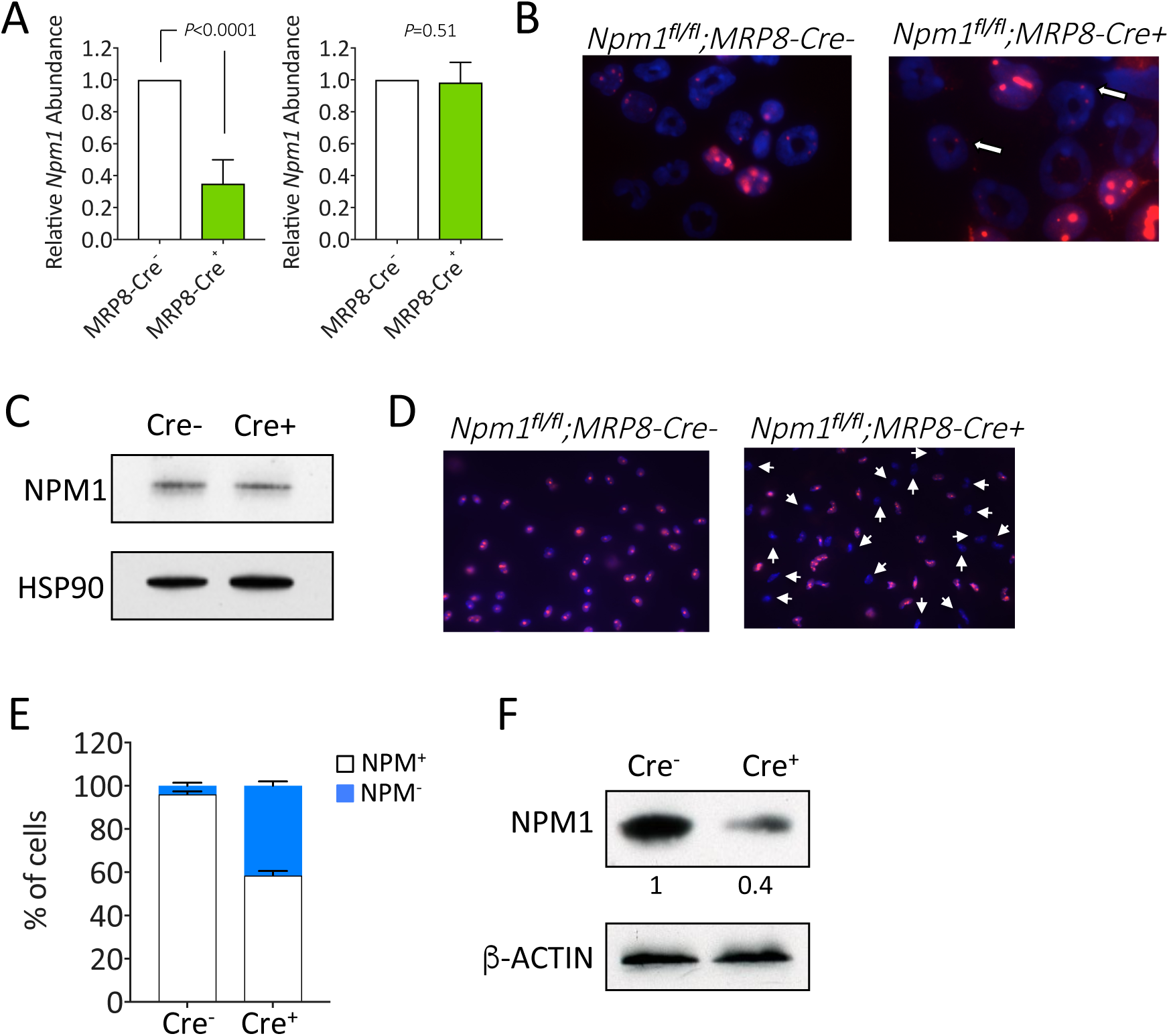
Npm1 loss in the myelo-monocytic lineage. (A) Genomic real time PCR on Gr1^+^Mac1^+^ and B220^+^ sorted bone marrow cells from *Npm1*^*fl/fl*^;*MRP8-Cre-* (white bars) and *Npm1*^*fl/fl*^;*MRP8-Cre*+ (green bars) mutant mice. (B) Immunofluorescence analysis of bone marrow cells from *Npm1*^*fl/fl*^;*MRP8-Cre*^*-*^ (left panel) and *Npm1*^*fl/fl*^;*MRP8-Cre*^+^ (right panel) using the monoclonal NPM antibody. White arrows indicate NPM positive nucleolar staining. (C) Western blot analysis on protein lysates from on Gr1^+^Mac1^+^ and B220^+^ sorted bone marrow cells using the monoclonal NPM1 antibody. (D) Immunofluorescence analysis of peritoneal macrophages from *Npm1*^*fl/fl*^;*MRP8-Cre*^*-*^ (left panel) and *Npm1*^*fl/fl*^;*MRP8-Cre*^+^ (right panel) using the monoclonal NPM antibody. White arrows indicate negative NPM nucleolar staining. (E) The percentage of NPM negative (white) and positive (blue) cells. (F) Western blot analysis on protein lysates of peritoneal macrophages from *Npm1*^*fl/fl*^;*MRP8-Cre*^*-*^ and *Npm1*^*fl/fl*^;*MRP8-Cre*^+^ using the monoclonal NPM1 antibody.

It is known that the NPM1 protein displays a very long half-life of approximately 36 hours(*8*), while the *in vivo* life-span of mature granulocytes is very short (<24 hours) (*9*). The long half-life of NPM1 might explain its presence in Gr1^+^;Mac1^+^ cells as the protein that was synthesized before genomic recombination remains present in the cell, despite the relatively efficient deletion of the gene in bone marrow granulocytes (Figure 1A).

Therefore, we chose to assess the efficiency of *Npm1* deletion by Cre recombinase on long-lived peritoneal macrophages. Thioglycolate induced peritoneal macrophages were isolated from *Npm1*^*fl/fl*^;*MRP8-Cre*^+^ and *Npm1*^*fl/fl*^;*MRP8-Cre*^*-*^ mice and plated on coverslips and immunofluorescence analysis for NPM1 was carried out over a 6-day time course. Almost 40% of peritoneal macrophages collected from *Npm1*^*fl/fl*^;*MRP8-Cre*^+^ mice displayed complete loss of *Npm1* by immunofluorescence analysis when compared with control macrophages (Figure 1D and 1E). Western blot analysis on protein lysates from cultured macrophages demonstrates a significant reduction of NPM1 in *Npm1*^*fl/fl*^;*MRP8-Cre*^+^ compared to *Npm1*^*fl/fl*^;*MRP8-Cre*^*-*^ cells (Figure 1F). Thus, these data demonstrate that *Npm1* can be efficiently knocked-out in the myeloid lineage, and does not appear to impact macrophage viability.

### Conditional inactivation of NPM1 in the myelomonocytic compartment did not impact blood counts and mice survival

We continued our study by analyzing *Npm1*^*fl/fl*^;*MRP8-Cre*^+^ mice for any alterations in blood count or in myeloid development. *Npm1*^*fl/fl*^;*MRP8-Cre*^+^ mice showed no major differences in complete blood counts of peripheral blood when compared to *Npm1*^*fl/fl*^;*MRP8-Cre*^*-*^ mice. Total white blood cell number (*Npm1*^*fl/fl*^;*MRP8-Cre*+: 3.7±0.98 x 10^3^/µL vs *Npm1*^*fl/fl*^;*MRP8-Cre*^*-*^ 4.3±1.48 *P*=0.71, Supplementary Figure 1A) granulocytes (*Npm1*^*fl/fl*^;*MRP8-Cre*+: 11.7±1.4% vs *Npm1*^*fl/fl*^;*MRP8-Cre-*9.9±1.9%, *P*=0.48, and Supplementary Figure 1B) and monocyte percentage (*Npm1*^*fl/fl*^;*MRP8-Cre*+: 1.43±0.14% vs *Npm1*^*fl/fl*^;*MRP8-Cre-*1.14±0.17% *P*=0.24, and Supplementary Figure 1C) were similar in both cohorts of mice (8-12 weeks of age). Morphological analysis on both granulocyte and monocyte cells in blood smears from these mice showed no signs of dysplasia at this young age (Supplementary Figure 1D). In addition, the overall survival of the *Npm1*^*fl/fl*^;*MRP8-Cre*^+^ mice did not appear to be affected, as a long term cohort (n=12) followed for a period of two years showed no overt disease (Supplementary Figure 1E).

### Npm1 deficient peritoneal macrophages display altered functions

As we demonstrated that NPM1 protein levels are reduced in long-lived peritoneal macrophages, we asked whether NPM1 loss may influence their survival and or function. In *ex vivo* culture, *Npm1* null macrophages were observed at the same numbers six days after plating, suggesting that NPM1 does not play a significant role in survival of these cells (data not shown). To determine whether NPM1 was able to impact macrophage effector functions, we measured respiratory burst and the production of reactive oxygen species (ROS), that are determinants of efficient killing by macrophages (*10*). *Npm1*^*fl/fl*^;*MRP8-Cre*^+^ peritoneal macrophages showed reduced superoxide production compared with *Npm1*^*fl/fl*^;*MRP8-Cre*^*-*^ macrophages (Figure 2A). Then, we determined the effect of *Npm1* deficiency on the ability of peritoneal macrophages to respond to chemo-attractants. Using the chemoattractant macrophage-inflammatory protein-2 (MIP2) *Npm1*^*fl/fl*^;*MRP8-Cre*^+^ and *Npm1*^*fl/fl*^;*MRP8-Cre*^*-*^ macrophages were evaluated for their ability to migrate through a transwell membrane. The chemotactic response of *Npm1*^*fl/fl*^;*MRP8-Cre*^+^ peritoneal macrophages was reduced by up to 46%±9.6 when compared to controls (Figure 2B). In order to test the effect of *Npm1* deletion on phagocytic activity, peritoneal macrophages were incubated with fluorescently labeled *Zymosan A* BioParticles. As shown in Figure 2C macrophages harvested from *Npm1*^*fl/fl*^;*MRP8-Cre*^+^ mice engulfed fewer particles than *Npm1*^*fl/fl*^;*MRP8-Cre*^*-*^ macrophages, indicating that they had a significantly reduced phagocytic activity.

**Figure 2.**
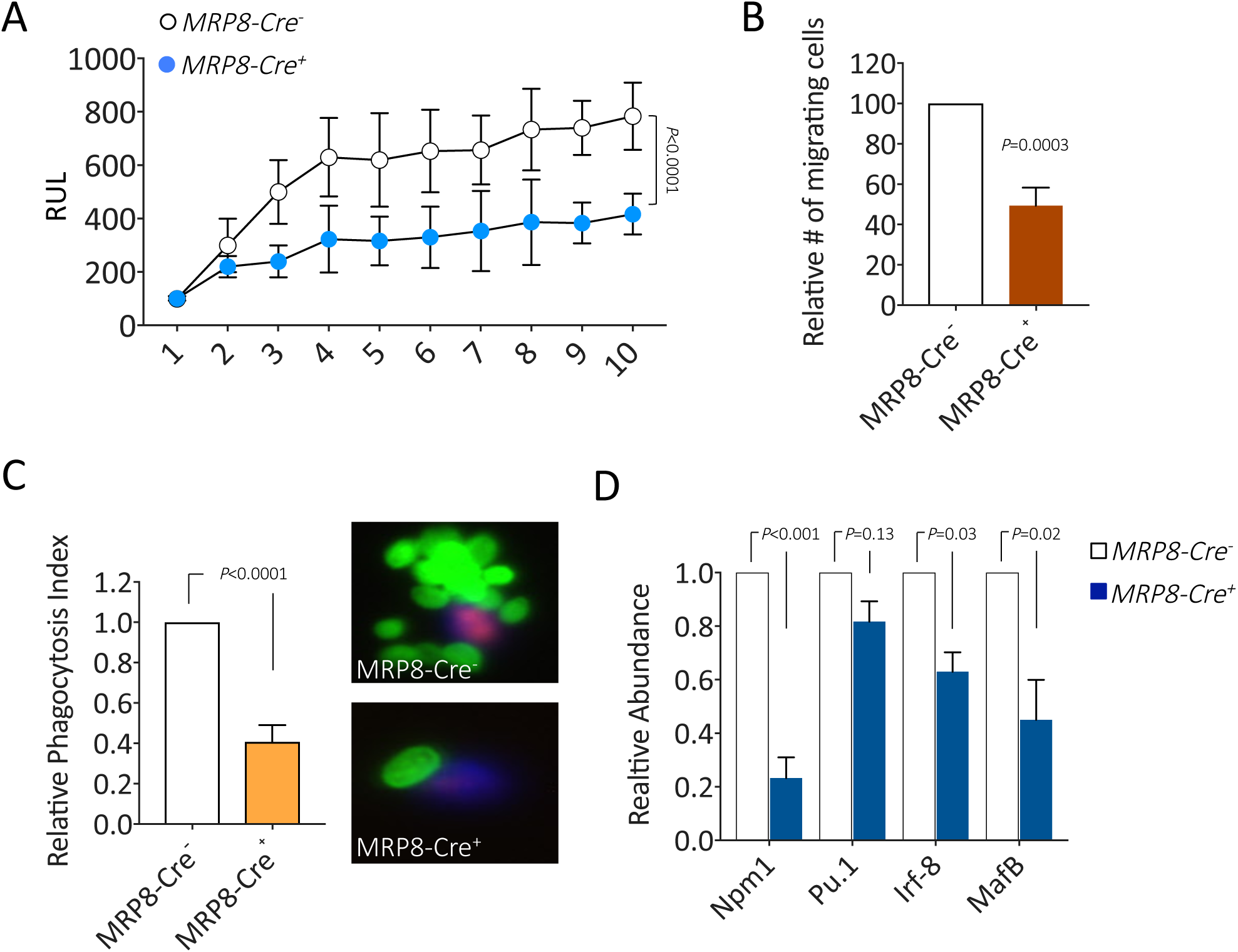
Npm1 deficiency leads to aberrant macrophage activity. (A) Measurement of ROS production by peritoneal macrophages from *Npm1*^*fl/fl*^;*MRP8-Cre*^*-*^ (white) and *Npm1*^*fl/fl*^;*MRP8-Cre*^+^(blue) mice after activation with phorbol myristate acetate. The results are displayed as the mean of triplicate samples±SD and analyzed by two way analysis of variance. (B) Chemotaxis of peritoneal macrophages from *Npm1*^*fl/fl*^;*MRP8-Cre*^*-*^ (white) and *Npm1*^*fl/fl*^;*MRP8-Cre*^+^ (red) mice in response to MIP2. The data represent mean±SD for three independent experiments. (C) Phagocytosis of Alexa Flour 488 conjugated zymosan A BioParticles by *Npm1*^*fl/fl*^;*MRP8-Cre*^*-*^ and *Npm1*^*fl/fl*^;*MRP8-Cre*^+^ peritoneal macrophages. Representative images of macrophages from *Npm1*^*fl/fl*^;*MRP8-Cre-* and *Npm1*^*fl/fl*^;*MRP8-Cre*+ mice. Results were presented as the average number of ingested beads per cell (phagocytic index = number of ingested beads/total number of macrophages). The data represent mean±SD for three independent experiments. (D) Relative expression levels of *Npm1, Pu.1, Irf-8* and *Maf-b* in *Npm1*^*fl/fl*^;*MRP8-Cre*^*-*^ (white bars) and *Npm1*^*fl/fl*^;*MRP8-Cre*^+^ (blue bars) macrophages as determined by quantitative RT-PCR analysis. The data represent mean±SD for three independent experiments.

Activated macrophages secrete multiple cytokines and chemokines that act via autocrine and paracrine pathways to initiate, amplify, and ultimately terminate the inflammatory response. Peritoneal macrophages, isolated from *Npm1*^*fl/fl*^;*MRP8-Cre*^+^ *and Npm1*^*fl/fl*^;*MRP8-Cre*^*-*^ mice, were stimulated with LPS. Compared with *Npm1*^*fl/fl*^;*MRP8-Cre-*macrophages, stimulation of *Npm1*^*fl/fl*^;*MRP8-Cre*^+^ macrophages with LPS did not modify the concentration of TNF or IL-6 production as measured by ELISA using culture supernatants (Supplementary Figure 2). Monocyte maturation toward macrophages is a finely regulated process that involves several transcription factors. Monocyte-lineage progression depends on the continued presence of PU.1, while Irf-8 may be an important PU.1 co-factor during monocyte maturation (*11*). Additionally, Maf-B is expressed at moderate levels in myeloblasts and is strongly up-regulated in both monocytes and macrophages (*12*). In order to establish whether the functional impairment shown by *Npm1* null macrophages would depend on an altered differentiation process we analyzed the expression of these key regulators of this process by qPCR. As shown in Figure 2D *Npm1*^*fl/fl*^;*MRP8-Cre*^+^ macrophages display a significant reduction of both *Maf-b* and *Irf-8* demonstrating a possible altered maturation process in absence of NPM1.

### NPM1-depletion leads to impaired function of human macrophage-like cells

Next, we wanted to understand whether the role of NPM1 in regulating macrophage function is conserved in human cells. To this end, we utilized the human monocyte cell line, THP-1, which can readily differentiate into macrophage-like cells upon stimulation with phorbol 12-myristate 13-acetate (PMA).

THP-1 cells were transduced with an shNPM1 lentiviral vector, which led to a very efficient depletion of NPM1 (figure 3A). Next, we performed functional analyses of differentiated control- and shNPM1-transduced THP1 cells. In accordance with the murine peritoneal macrophages data, we found that differentiated NPM1-depleted THP1 cells show reduced functional capacities. Specifically, we found that upon re-stimulation NPM1-depleted cells produce significantly lower amount of ROS (Figure 3B). Additionally, NPM1-depleted cells also demonstrated reduced phagocytosis of labeled particles as compared to control cells (Figure 3D). Interestingly, when cytokine production was tested, we observed reduced levels of IL-β and IL-6 2.5hrs after stimulation (but not of TNF, Figure 3D), and no differences in cytokine levels 24hrs after stimulation (Supplementary Figure 3). This response curve might explain why no differences were observed in the murine peritoneal macrophages cytokine secretion capacity, as it was tested only 24 hrs after LPS stimulation. Taken together, these data demonstrate that NPM1 has a conserved role in facilitating macrophage functions in both human and mouse.

**Figure 3.**
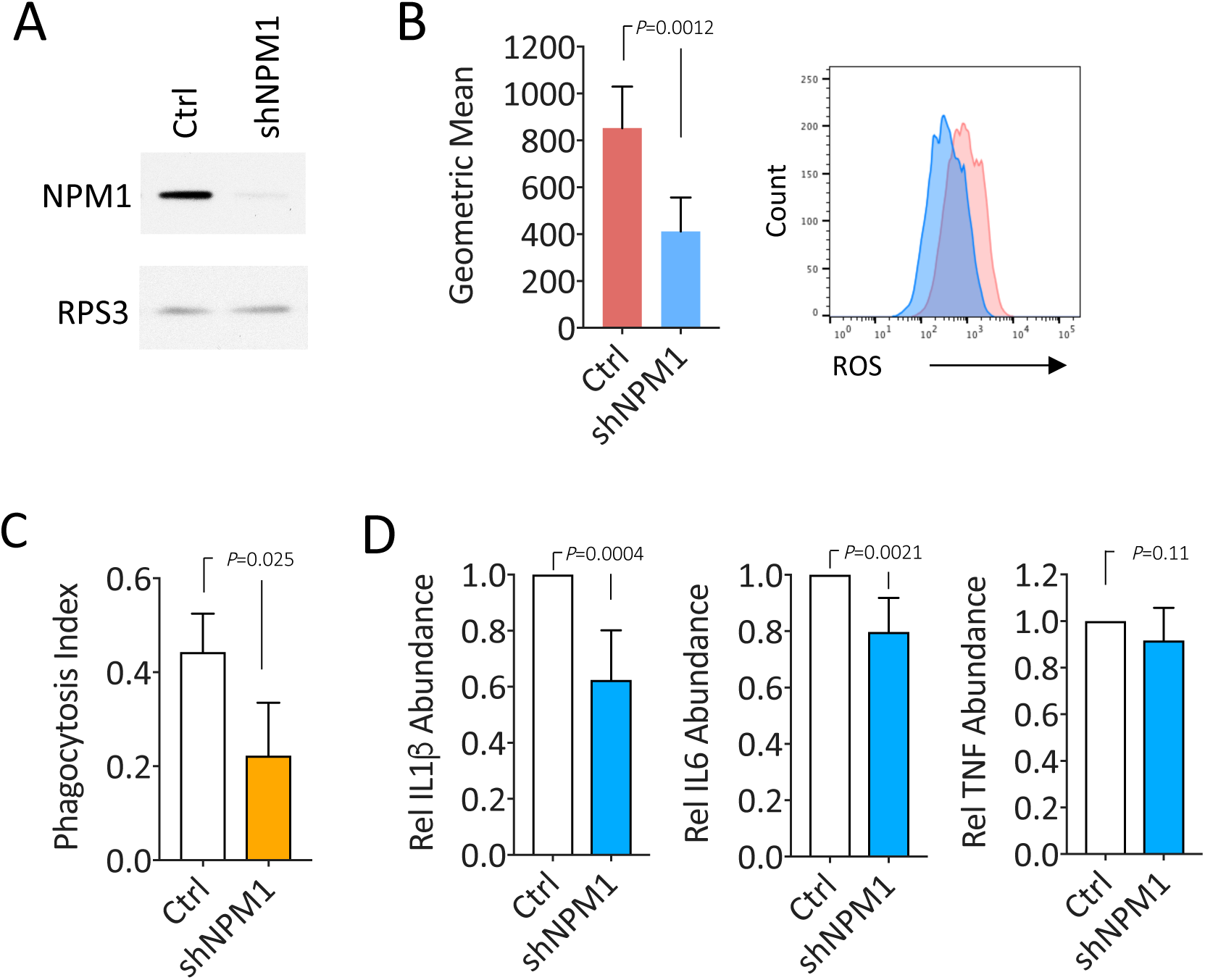
Human macrophage-like cells show reduce activity upon NPM1 depletion. (A) Western blot of THP1 cells transduced with anti-NPM1 shRNA lentivirus. (B) Measurement of ROS production by PMA-differentiated THP1 cells transduced with the indicated lentivirus. ROS measurement was performed 30 minutes after PMA re-stimulation. Left - data represent Geometric Mean±SD for three independent experiments. Right – a representative example of ROS production. (C) Phagocytosis of BioParticles by either control or shNPM1 PMA-differentiated THP1 cells. (D) Quantitative real time PCR analysis of the relative abundance of ILβ, IL6 and TNF in PMA-differentiated THP1 cells transduced either with control vector or shNPM1. Data are Mean±SD for three independent experiments.

### NPM1 controls macrophage function partially through regulation of ribosome function

We recently demonstrated the role of NPM1 as an RNA binding protein that mediates 2’-O-methylation of ribosomal RNA (rRNA) through its ability to bind C/D-box snoRNA (*3*). Through this function it may contribute to ribosome cell-specific function and regulate translation. In this study we also identified a novel germline mutation in NPM1 in a bone marrow failure patient, *NPM1*^*D180del*^, that specifically diminishes its ability to bind C/D-box snoRNAs and regulate translation. We generated a knock-in mouse model of the *NPM1*^*D180del*^ mutation and demonstrated that these mice suffer from bone marrow failure, causally linking regulation of 2’-O-methylation and ribosome function to hematological disease.

In order to gain more molecular insight into the mechanism by which NPM1 controls macrophage function, and to understand to which extent the ribosome and translation play a role in this, we analyzed the functional capacity of *NPM1*^*D180del*^ derived peritoneal macrophages. In contrast to *Npm1*^*f/f*^;*MRP8-Cre*^+^ macrophages, we found *NPM1*^*D180del*^ macrophages show neither differences in their capacity to produce ROS upon LPS stimulation (Figure 4A), nor in their migration in response to a chemoattractant (Figure 4B). Most importantly, however, and in accordance with *Npm1*^*f/f*^;*MRP8-Cre*^+^ macrophages as well with shNPM1 THP1 cells, *NPM1*^*D180del*^ macrophages exhibit reduced phagocytosis of labeled particles (Figure 4C) as well as lower levels of IL-β and IL-6, but not of TNF (Figure 4D). These data demonstrate that NPM1 controls macrophage function in multiple ways, including through regulation of translation.

**Figure 4.**
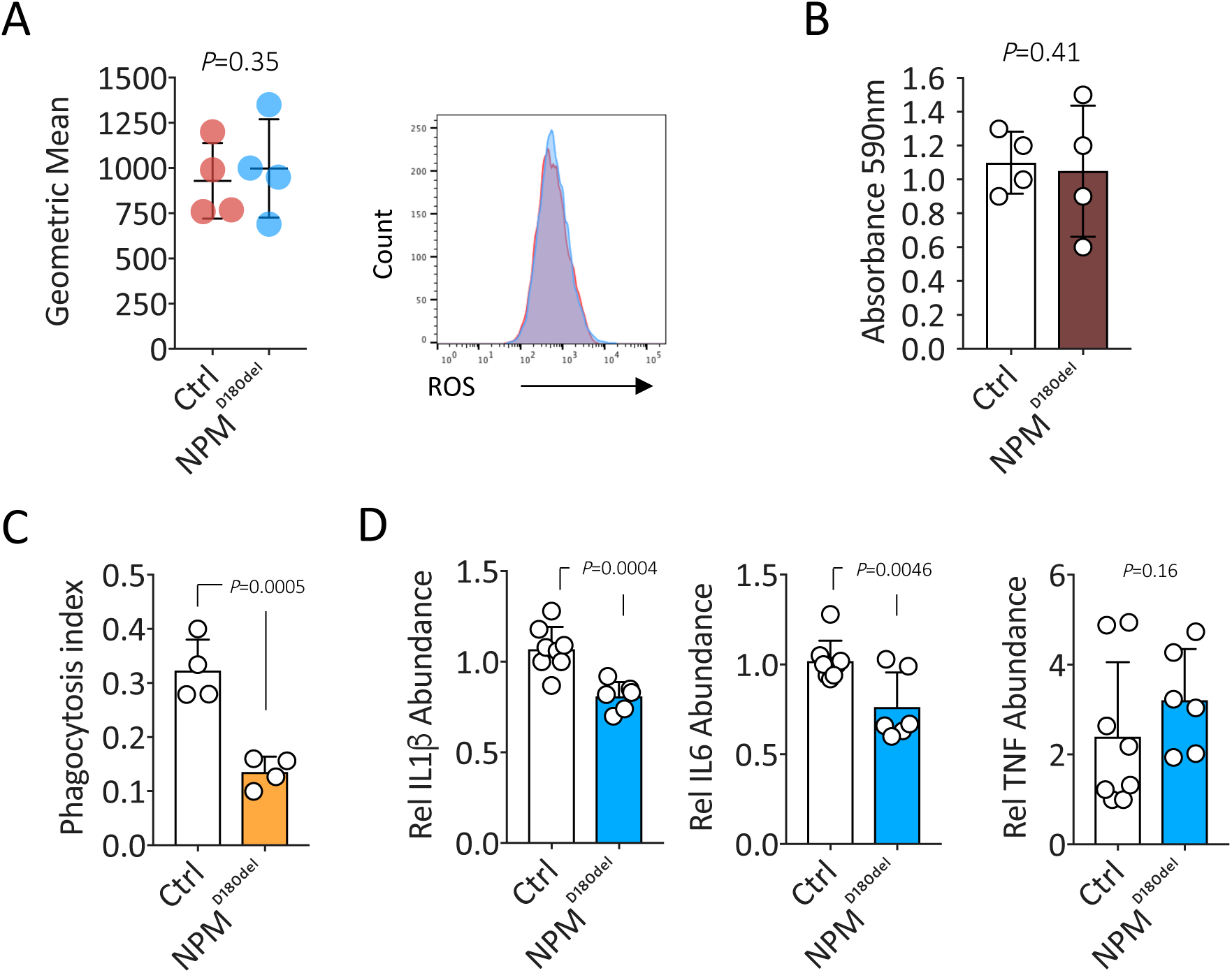
Ribosome dysfunction leads to aberrant macrophage function. (A) Measurement of ROS production by *NPM*^*D180del*^ (blue) and wild type littermates (red) peritoneal macrophages. Left - data represent Geometric Mean±SD for four independent experiments. Right – a representative example of ROS production. (B) Chemotaxis of peritoneal macrophages from *NPM*^*D180del*^ (red bar) and wild type littermates (white bar) mice in response to MIP2. The data represent mean±SD for four independent experiments. (C) Phagocytosis of BioParticles by *NPM*^*D180del*^ (orange bar) and wild type littermates (white bar) mice peritoneal macrophages. Results were presented as the average number of ingested beads per cell (phagocytic index = number of ingested beads/total number of macrophages). The data represent mean±SD for four independent experiments. (D) Quantitative real time PCR analysis of the relative abundance of ILβ, IL6 and TNF in *NPM*^*D180del*^ (blue bar) and wild type littermates (white bar) mice peritoneal macrophages, 2.5hrs after stimulation with LPS (100ng/ml). Data are Mean±SD for six independent experiments.

## Discussion

NPM1 is a shuttling protein that interacts with multiple partners involved in several different cellular functions (*1*). The *NPM1* gene undergoes frequent genetic alterations in hematological tumors (*1*). These alterations include deletions in MDS, chromosomal translocations such as with the *ALK* gene in anaplastic large cell lymphoma, with the *MLF1* and *RARA* genes in AML and acute promyelocytic leukemia, respectively(*1,13-16*). Additionally, *NPM1* bears the most frequent mutation in AML with normal karyotype, as approximately 30% of patients present with a frame-shift mutation in NPM1 (*4*). However, even though NPM1 is so intensively studied, its function in mature adult myelopoiesis and in cellular function is so far not well defined. For this reason, we have generated specific *Npm1* conditional mutants in the hematopoietic system, using the *Mrp8-Cre* model, a model that allows conditional inactivation of *Npm1* across myelo-monocytic compartment. Using this genetic mouse model we functionally ablated NPM1 across myelomonocytic lineage. This allowed us to demonstrate that while altered *NPM1* dosage does not affect myeloid development, it does disrupt macrophage effector capacity, by inhibiting chemotactic, phagocytic functions and ROS production, that are critical functions of macrophages.

Interestingly, a previous report suggested enhanced, rather than reduced, macrophage maturation and function in *Npm1* heterozygous mice (*17*). These presumed discrepancies might arise due to various reasons. First, different mice models were used in the two studies. Guery et al. used a total body heterozygous model of *Npm1*, affecting the hematopoietic system in an unspecific manner, and possibly affecting additional organs and tissues. However, in the current study we used a conditional deletion of *Npm1* specifically in the myelomonocytic lineage, thereby minimizing cell extrinsic effects on macrophage function. In addition, we further demonstrate the functional role of NPM1 in additional models such as human cells and another genetic mouse model of a specific *Npm1* mutation. Second, Guery et al, performed their study in heterozygous setting, while we performed our study under complete inactivation of *Npm1*. Future studies will reveal which of the functions of NPM1 are loss due to heterozygosity, and which are abolished only after complete loss of NPM1, and thus will shed light on the mechanisms and pathways that control macrophage activity.

It is becoming evident that translation regulation has a major role in many biological processes (*18-20*), however its role in immune function is poorly documented. Here we report that alterations to ribosomal modifications can influence macrophage function. A mouse model with a specific mutation in *Npm1* with impaired ability to bind C/D-box snoRNAs and properly methylate ribosomal RNA results in altered cytokine levels and phagocytosis. Thus, we establish a central role for ribosome regulation in mediating macrophage function. It is becoming clear that active translation regulation by the ribosome, either through changes to ribosome composition (*20-22*) or to ribosome numbers (*19,23*), is an important and novel layer of gene expression regulation. Thus, future studies are warranted to better understand the centrality of the ribosome in the regulation of immune function.

## Acknowledgements

This work was funded by National Institutes of Health grants, CA-71692 and CA-74031, and by an Outstanding Investigator Award R35 (CA197529) grant and the SHINE (5R01DK115536) grant to P.P.P.. P.S. was supported in part by the Associazione Umbra Leucemie e Linfomi (AULL).

## Authorship Contribution

D.N. and P.S. designed and performed experiments, analyzed data and wrote the paper; J.G.C. performed experiments and wrote the paper; L.R. performed experiments; A. K-G. and A.M. analyzed data and wrote the paper; J.C. performed experiments and analyzed data; N.B. analyzed data and wrote the paper; P.P.P. designed research, analyzed data and wrote the paper.

## Conflict-of-interest disclosure

The authors declare no competing financial interests.

## Methods

### Isolation of peritoneal macrophages

Thioglycolate-elicited murine peritoneal macrophages were harvested 5 days after i.p. injection of 1 ml of 2.9% Brewer’s complete thioglycolate broth. Cells were plated and only adherent cells were used in experiments.

### Superoxide production and chemotaxis assays

Assays for the respiratory burst and chemotaxis were performed using luminol-enhanced chemiluminescence (Diogenes, National Diagnostics, Atlanta, GA) in cells activated using phorbol myristate acetate (PMA, 3.2 µM, Sigma). ROS production, by differentiated THP1 cells and peritoneal macrophages derived from *Npm1*^*D180del*^ mice, was measured by CellROX deep red flow cytometry assay. Chemotaxis assays were performed using Corning Transwell 24 well plates with 3.0 um pore polycarbonate membranes (Corning Life Sciences, Acton, MA), and the mouse chemoattractant macrophage-inflammatory protein-2 (MIP-2, R&D Systems). Migration of cells was evaluated by trypan blue exclusion following incubation.

### Phagocytosis assay

Phagocytosis was measured using Alexa Flour 488 conjugated zymosan A BioParticles (Molecular Probes z-23373) along with a specific opsonizing reagent (Molecular Probes z-2850). Zymosan A BioParticles and opsonizing reagents were reconstituted according to the manufacturers instruction. Peritoneal macrophages cultured in 60 mm dishes containing coverslips were allowed to ingest zymosan particles for 1 h at 37°C. Non-ingested particles were removed by three washes in PBS. For *Npmfl/fl;MRP8-Cre* derived peritoneal macrophages – cells were fixated with 4% PFA and then stained with anti-NPM1 antibody in order to distinguish NPM1 negative and positive cells. Phagocytosed particles were counted using a fluorescent microscope and the average number of particles per macrophage was calculated after viewing 100 cells in 3 different fields. Free zymosan BioParticles were easily distinguished from cells containing zymosan by the presence within the cytoplasmic membrane in phase microscopy.

### Cytokine quantification

Cells were cultured overnight in the presence of 100 ng/ml LPS. Cell supernatants were harvested, centrifuged to remove dead cells, and analyzed by ELISA using cytokine-specific Quantikine ELISA kits from R&D Systems (Minneapolis, MN).

### Statistical analysis

Data were analyzed using a paired Student’s t test and the analysis of variance (ANOVA).

**Supplementary Figure 1.**
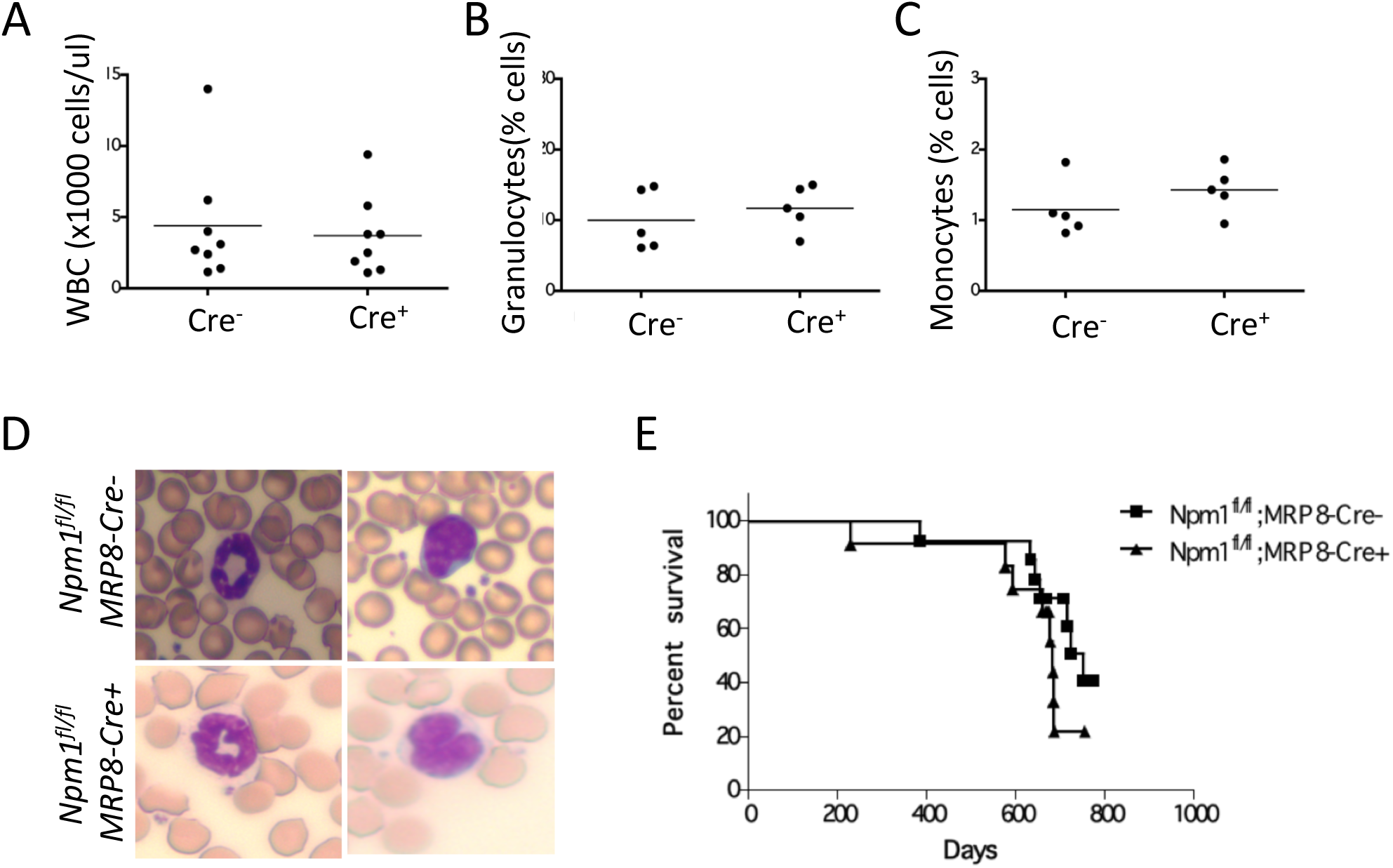
(A) Peripheral white blood cells count of *Npm1*^*fl/fl*^;*MRP8-Cre*^*-*^ and *Npm1*^*fl/fl*^;*MRP8-Cre*^+^ mice. (B) and (C) Percentages of granulocytic and monocytic cells (respectively) in the peripheral blood of *Npm1*^*fl/fl*^;*MRP8-Cre*^*-*^ and *Npm1*^*fl/fl*^;*MRP8-Cre*^+^ mice. (D) 60X magnification of Giemsa stained peripheral blood smears from *Npm1*^*fl/fl*^;*MRP8-Cre-* (upper panels) and *Npm1*^*fl/fl*^;*MRP8-Cre*+ (lower panels) mice showing no major differences in granulocytes (left panels) and monocytes (right panels) morphology. (E) Kaplan-Meier survival curve demonstrating no significant differences regarding overall survival between *Npm1*^*fl/fl*^;*MRP8-Cre-* (n=12) and *Npm1*^*fl/fl*^;*MRP8-Cre*+ mice (n=12).

**Supplementary Figure 2.**
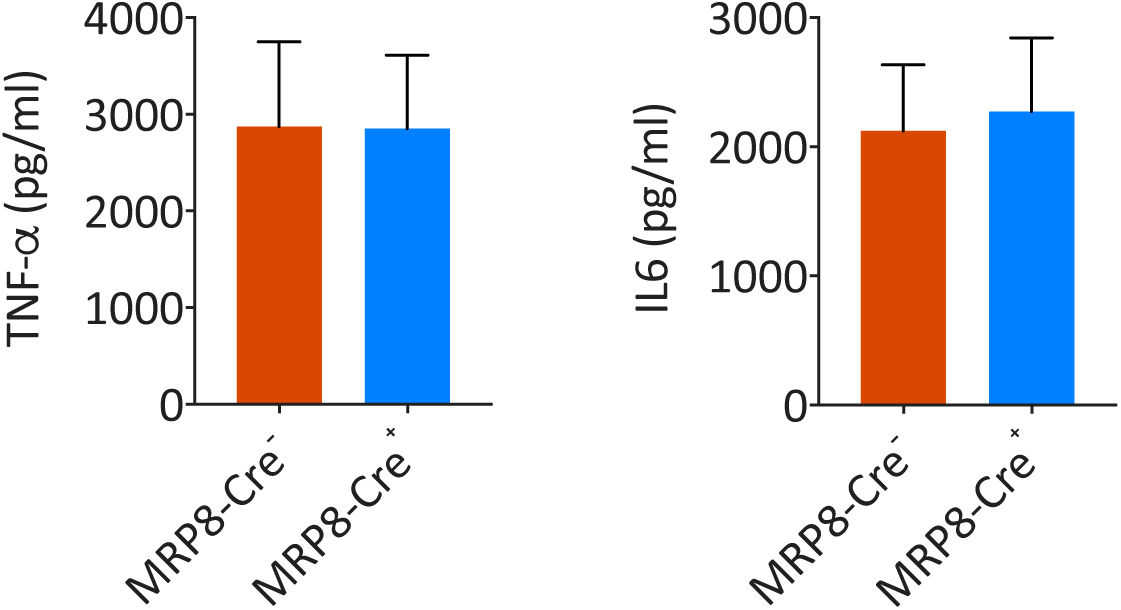
Release of TNF-α (left graph) and IL-6 (right graph) cytokines from peritoneal macrophages from *Npm1*^*fl/fl*^;*MRP8-Cre-* (red bar) and *Npm1*^*fl/fl*^;*MRP8-Cre*+ (blue bar) mice after stimulation with LPS. The data represent mean±SD for five independent experiments.

**Supplementary Figure 3.**
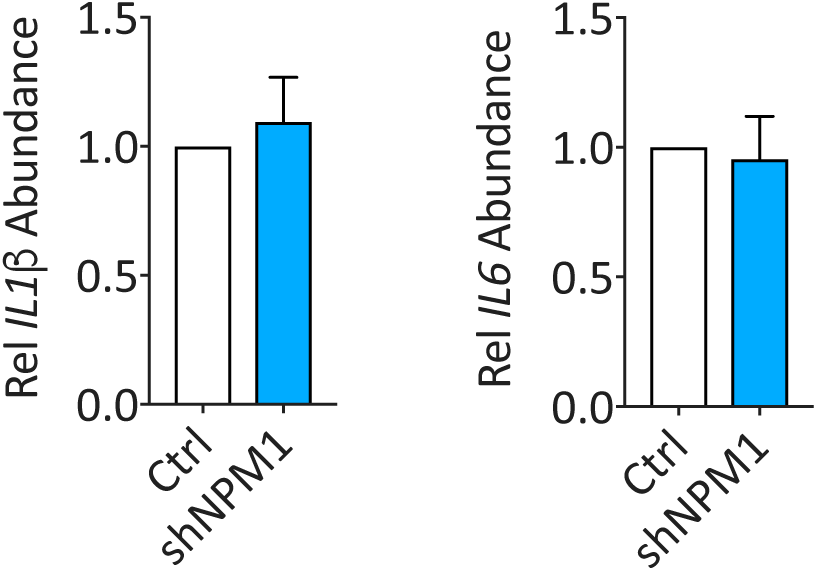
Quantitative real time PCR analysis of the relative abundance of ILβ and IL6 in control (white bar) and shNPM1 (blue bar) PMA-differentiated THP1 cells, 24hrs after stimulation with PMA. Data are Mean±SD for three independent experiments.

## Notes

### Competing Interest Statement

The authors have declared no competing interest.

